# Pathogenic Mutations in the C2A Domain of Dysferlin form Amyloid that Activates the Inflammasome

**DOI:** 10.1101/2023.04.24.538129

**Authors:** Isaac L. Scott, Matthew J. Dominguez, Adam Snow, Faraz M. Harsini, Justin Williams, Kerry L. Fuson, Roshan Thapa, Pushpak Bhattacharjee, Gail A. Cornwall, Peter A. Keyel, R. Bryan Sutton

## Abstract

Limb-Girdle Muscular Dystrophy Type-2B/2R is caused by mutations in the *dysferlin* gene (*DYSF*). This disease has two known pathogenic missense mutations that occur within dysferlin’s C2A domain, namely C2A^W52R^ and C2A^V67D^. Yet, the etiological rationale to explain the disease linkage for these two mutations is still unclear. In this study, we have presented evidence from biophysical, computational, and immunological experiments which suggest that these missense mutations interfere with dysferlin’s ability to repair cells. The failure of C2A^W52R^ and C2A^V67D^ to initiate membrane repair arises from their propensity to form stable amyloid. The misfolding of the C2A domain caused by either mutation exposes β-strands, which are predicted to nucleate classical amyloid structures. When dysferlin C2A amyloid is formed, it triggers the NLRP3 inflammasome, leading to the secretion of inflammatory cytokines, including IL-1β. The present study suggests that the muscle dysfunction and inflammation evident in Limb-Girdle Muscular Dystrophy types-2B/2R, specifically in cases involving C2A^W52R^ and C2A^V67D^, as well as other C2 domain mutations with considerable hydrophobic core involvement, may be attributed to this mechanism.

## Introduction

The large mechanical forces generated by muscle cells cause microtears within their membrane [40]. The propensity for muscle tissue to develop microtears necessitates an efficient, rapid *in vivo* repair mechanism to ensure muscle tissue function and survival [35, 41]. Multiple proteins have been implicated in skeletal muscle membrane repair; however, the dysferlin protein has been implicated in the membrane targeting and membrane fusion steps of the repair process [2, 9]. Mutations within the *DYSF* gene cause Limb-Girdle Muscular Dystrophy Type-2B/2R (LGMD) and Miyoshi Myopathy (MM) in humans. Both conditions are typically diagnosed in late childhood and lead to loss of independent ambulation late in adulthood, with severe cases compromising cardiac and respiratory function [20, 64].

Wild-type dysferlin is a 237 kDa single transmembrane-spanning protein with multiple tandem C2 domains and a unique Arg/Trp-rich DysF domain. The dysferlin primary sequence has historically been annotated with as many as seven C2 domains; each C2 domain has been assigned a letter (A-G), signifying the relative order of each domain in the protein’s primary sequence [20]. More recently, an analysis of ferlin domain structure revealed an additional C2 domain between C2C and C2D. As the eighth C2 domain subsumes the four-helix bundle FerA domain of dysferlin, it was designated as C2-FerA [12].

The current hypothesis that describes dysferlin’s physiological activity is the ‘patch repair’ model [18]. In this model, the C2 domains of dysferlin act as sensors to detect an influx of exogenous Ca^2+^ that signals damage within the muscle cell membrane. Once localized to the site of damage, dysferlin mediates Ca^2+^-dependent vesicle aggregation. The aggregated vesicles then form a patch that fuses with the cell membrane to repair the site of damage [8, 15, 50].

More than 1300 distinct pathogenic variants within *DYSF* transcripts have been identified to date [6]. Most are nonsense mutations that truncate the dysferlin protein; however, approximately 270 are missense mutations. Among the set of pathogenic missense mutations known in *DYSF*, two clinically-described mutations occur in the first C2 domain of dysferlin (C2A), C2A^W52R^ (Trp-52 to Arg) and C2A^V67D^ (Val-67 to Asp) (Figure 1). The C2A^W52R^ was described in LGMD patients whose serum creatine kinase (CK) levels were 50 times normal [34, 37]. C2A^V67D^ was first described as a homozygous mutation of consanguineous origin in a large Russian family wherein affected individuals were diagnosed with either distal myopathy or LGMD-2B/2R [27]. Affected individuals begin to develop characteristic muscle weakness between 15 and 20 years of age [27]. Western blot analyses from muscle biopsies revealed that clinical tissue samples possessed no detectable amounts of dysferlin, suggesting the protein may mislocalize [1, 52, 57]. In addition to mislocalizing, dysferlin C2A^W52R^ and C2A^V67D^ mutations alter interactions with MG53 in pull-down assays [39]. Similarly, C2A^V67D^ fails to bind to AHNAK in pull-down assays [26]. *In vitro*, Dysferlin C2A^V67D^ has reduced phospholipid-binding activity, and an inability to bind to membranes [10, 25].

**Figure 1.**
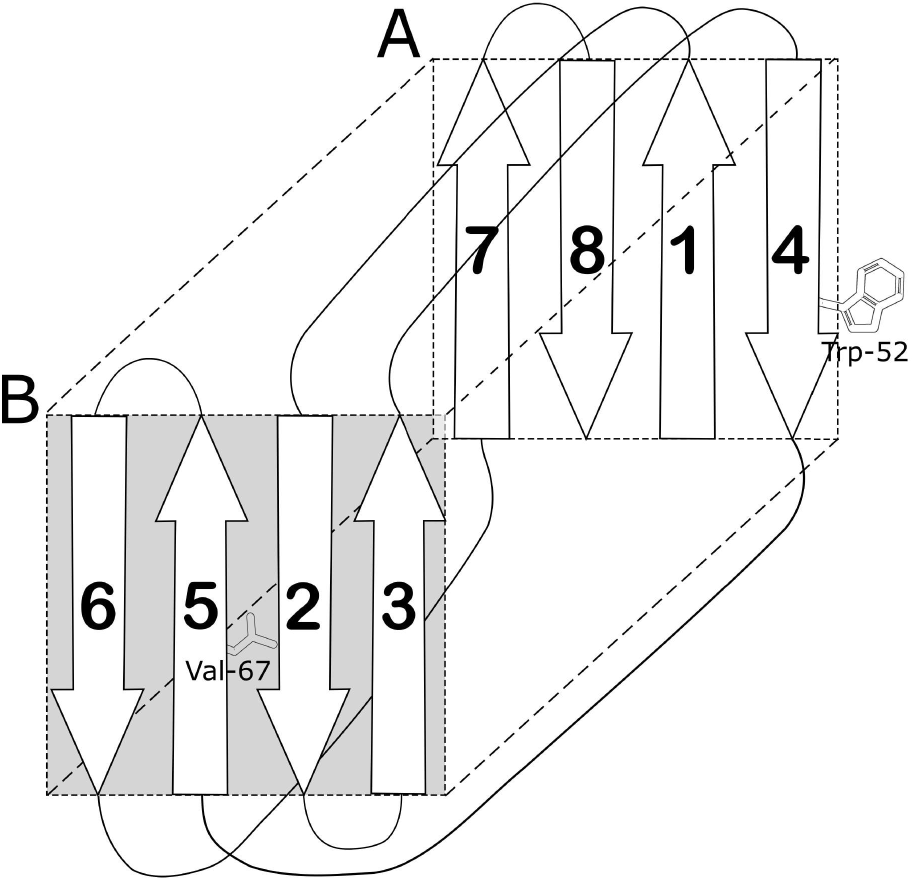
Schematic of C2 domain of dysferlin C2A. Sheets A and B are highlighted.

The previous data compiled for these two mutations support a severe dysferlin protein defect. However, the mechanism by which a single-point mutation in one domain alters the activity of a 237-kDa protein remains unknown. Given the dysferlin’s highly modular architecture, we used biophysical analysis, molecular dynamics, and cell biology to determine the mechanistic impact of the mutations in both the isolated C2A domain and the whole protein. We found that the dysferlin C2A^W52R^ and the C2A^V67D^ mutations can disrupt the typical hydrogen-bonded β-sheet structure of the C2A domain, which exposes sequences predicted to be amyloidogenic. The mutated dysferlin C2A domains form a cross β-amyloid structure that impacts the membrane repair capacity of the entire dysferlin molecule. Furthermore, the resulting amyloid stimulates the inflammasome to secrete pro-inflammatory proteins like IL-1β. These data provide experimental evidence for a link between dysferlin mutations and the chronic inflammatory state observed in dystrophic patients’ muscle tissue [4, 47, 63].

## Materials and Methods

### Reagents

All reagents were from Thermo Fisher (Waltham, MA, USA) unless otherwise specified. Nigericin and propidium iodide (PI) (Lot MKCB0899V) was from Sigma-Aldrich (St. Louis, MO, USA). 3,4-methylenedioxy-β-nitrostyrene (MNS) was from Tocris Biosciences (Minneapolis, MN), while ultrapure LPS from *Escherichia coli*, 0111:B4 strain and monosodium urate (MSU) were from Invivogen (San Diego, CA). Anti-mouse IL-1β ELISA capture antibody B122 (Catalog 14-7012-85, Lot E05277-1632) and biotinylated rabbit anti-mouse IL-1β detection polyclonal antibody (Catalog 13-7112-85, Lot 4285171) were from eBioscience (San Diego, CA). TMB and Streptavidin conjugated to horseradish peroxidase (HRP) (Catalog 405103, Lot B167931) were from Biolegend (San Diego, CA). Pierce lactate dehydrogenase (LDH) Cytotoxicity Assay Kit was from Thermo Fisher Scientific.

### Plasmids

The pET22b plasmids encoding His-tagged aerolysin and aerolysin^Y221G^ were kind gifts from Gisou van der Goot (Ećole Polytechnique Fédérale de Lausanne, Canton of Vaud, Switzerland) [60]. The wild-type aerolysin contained Q254E, R260A, R449A, and E450Q mutations, and the C-terminal KSASA was replaced with NVSLSVTPAANQLE HHHHHH compared to sequences in GenBank (i.e. M16495.1). The pBAD-gIII plasmid encoding His-tagged codon-optimized SLO was previously described [48]. The GFP-dysferlin plasmid was previously described [58]. The C2A^W52R^ and C2A^V67D^ mutations were introduced into GFP-dysferlin using Q5 mutagenesis (New England Biolabs).

### Purification of Dysferlin C2A domains and preparation of dysferlin C2A amyloid

Wild-type Human Dysferlin C2A protein was prepared as previously described [17]. Due to the propensity of C2A^W52R^ and C2A^V67D^ to aggregate during purification, the mutant C2A domains were purified using a modified method. Genes coding for human dysferlin C2A^W52R^ and C2A^V67D^ were cloned into the pGEX-4T vector plasmid. The resultant recombinant plasmids were transformed into ultra-competent ArcticExpress (DE3) cells (Agilent Technologies). ArcticExpress cells are derived from BL21(DE3), but they co-express two cold-adapted chaperonins that aid in folding proteins at low temperatures. Six liters of ArcticExpress-containing cells were grown in TB media at 37 °C until the cell density reached an OD_600_ of 1. The culture was then cooled to 10 °C and 400 µL of 1 M IPTG was added to initiate heterologous protein expression. The cultures were grown overnight at 10 °C while shaking at 250 rpm. Cells were harvested via centrifugation, pellets were flash-frozen in liquid Nitrogen, and stored at -80 °C.

For C2A^W52R^ and C2A^V67D^ purification, cells stored at -80 °C were thawed on ice. Cells were then resuspended in chilled wash buffer (40 mM HEPES, 150 mM NaCl, 1 mM CaCl_2_, pH 7.4, filter sterilized (.22µm)) and lysed via M110-EH Microfluidizer. The lysate was centrifuged in a JA-20 rotor for 45 minutes at 19.5k RPM (46k x g). The clarified supernatant was decanted, pooled, and combined with glutathione-S-Transferase (GST) resin. The resin and lysate slurry was poured into a column and allowed to settle under gravity for 2 hrs on ice. Flow-through was collected separately. The column was then washed with 20 column volumes of chilled wash buffer. The mutant C2A domains were digested with human α-thrombin on the column (5 µL at 5.7 mg/mL (21.15 units)) in 20 mL chilled wash buffer. The thrombin solution was applied to resin via stirring and then allowed to incubate overnight at 4 °C. After incubation, the eluate was collected, the thrombin was inactivated with 0.5 mM phenylmethylsulfonyl fluoride (PMSF), and the column bed was washed out with 10 mL chilled wash buffer (Figure S1). Purified C2A^W52R^ and C2A^V67D^ domains were stripped of bound Ca^2+^ with 1 mM EGTA to initiate amyloid formation.

### Recombinant toxins

Pore-forming toxins were expressed in *E. coli* and purified as previously described [33, 49, 58]. Briefly, toxin expression was induced with 0.2% arabinose for SLO, or 0.2 mM IPTG for aerolysin for 3 h at room temperature and then purified using Nickel-NTA resin. The protein concentration was determined by Bradford Assay. The hemolytic activity of each toxin was determined as previously described [33, 49] using human red blood cells (Zen-Bio, Research Triangle Park, NC, USA). One hemolytic unit (HU) is defined as the quantity of toxin required to lyse 50% of a 2% human red blood cell solution in 30 min at 37°C in 2 mM CaCl_2_, 10 mM HEPES (pH 7.4) and 0.3% BSA in PBS [32, 49]. We used units of HU/mL to normalize toxin activities in each experiment and to achieve consistent cytotoxicity across toxin preparations. The specific activity of toxins was also determined (Table 1)

**Table 1.** Specific activities of the toxins used in this study. N/D (not determined)

| Toxin | Hemolytic Activity (HU/mL) | Protein Conc (mg/mL) | Specific Activity (HU/mg) |
| --- | --- | --- | --- |
| aerolysin | 5x 10<sup>5</sup> | 2.8 | 1.8x 10<sup>5</sup> |
| SLO | 4x 10<sup>5</sup> | 2.4 | 1.7x 10<sup>5</sup> |
| aerolysin<sup>Y221G</sup> | <1000 | 1.6 | N/D |

### Mice

All experimental mice were housed and maintained according to Texas Tech University Institutional Animal Care and Use Committee (TTU IACUC) standards, adhering to the Guide for the Care and Use of Laboratory Animals (8th edition, NRC 2011). TTU IACUC approved mouse use. Bone marrow was obtained from C57BL/6 (B6), NLRP3^-/-^ or Casp1/11^-/-^ mice on a B6 background (Jackson Labs, Bar Harbor, ME, stocks 000664, 021302, 016621, respectively) as previously described [24, 29]. Mice of both sexes aged 6–15 weeks were used to prepare BMDM (Bone Marrow-derived Macrophages). The sample size was determined as the minimum number of mice needed to provide enough bone marrow for experiments. Consequently, no randomization or blinding was needed. Mice were euthanized by asphyxiation through the controlled flow of pure CO_2_ followed by cervical dislocation.

BMDM were prepared as previously described [24, 29]. Briefly, bone marrow was flushed from femora and tibiae, and the cells were cultured at 37 °C and 5% CO_2_ for 7–21 days on bacterial grade 15 cm tissue culture plates in 20% FBS (Premium grade, Seradigm (Radnor, PA)), 1 mM sodium pyruvate (Corning, Manassas, VA), 2 mM L-glutamine, 100 U/mL penicillin/100 µg/mL streptomycin (HyClone) and 30% L cell supernatant. Media was replaced on day 4. The cells were harvested using Cellstripper (Corning, Manassas, VA) and plated in Iscove’s modified Dulbecco’s medium (Corning, Manassas, VA) supplemented with 10% FCS, 2 mM L-glutamine and 100 U/mL penicillin /100 µg/mL streptomycin one day prior to experiments. L cell supernatant was media conditioned by L929 cells (American Type Culture Collection, catalog CCL-1) for 6 days as previously described [24].

### Cell culture

All cells were maintained at 37 °C, 5% CO_2_ and were negative for mycoplasma. HeLa (Catalog CCL-2, ATCC, Manassas, VA, USA) were cultured in DMEM (Corning, Corning, NY, USA) supplemented with 10% fetal calf serum (FCS) (Atlas Biologicals, Fort Collins, CO, USA) and 1x L-glutamine (Hyclone, Logan, UT).

### Negative stain electron microscopy

After the addition of 1 mM EGTA to initiate amyloid formation, five µl of purified C2A^WT^, C2A^W52R^, or C2A^V67D^ was spotted for one minute on 200 mesh nickel grids coated in formvar/carbon (Ted Pella, Redding, CA). The sample was blotted with filter paper, washed in water for 1 minute, stained for 1 minute with 2 percent uranyl acetate, and then washed in water again for 1 minute. A Hitachi 8100 electron microscope was used to analyze the samples, using an excitation voltage of 75 kV.

### Circular Dichroism Spectroscopy

30 µM of purified (C2A^WT^, C2A^W52R^ or C2A^V67D^) was diluted into 5 mM Tris, pH 7.4, and either 1 mM CaCl_2_ or 1 mM EGTA and pipetted into a 0.1 cm quartz cuvette. Spectra for the mutant C2A domains were measured immediately after elution from the GST column, and prior to amyloid formation. CD spectra were collected on a Jasco J-815 Circular Dichroism (CD) Spectropolarimeter at 20 °C. Measurements were collected from the wavelength range 200-250 nm for each protein domain (Figure S2). Ten accumulations were collected and averaged, with each sample measured at least three times. The secondary structure was assessed by BestSel using a common wavelength range from 200 - 250 nm [43].

### ThT dye binding assay

Samples of either C2A^WT^, C2A^W52R^ or C2A^V67D^ were filter sterilized (0.22 µm) and adjusted to 1 mg/mL. Ten µL of purified C2A domain (10 µg) were combined with 20 µL 1 mM ThT, + PBS to 100 µL total volume. Some samples included 2 µL 500 mM EGTA. All three domains were run in triplicate, and ThT controls were run in duplicate. All readings were performed at 37 °C in a black flat-bottom, 96-well plate. Buffer controls included all assay components in the absence of protein. Fluorescence was measured using a Tecan plate reader (excitation 450 nm, emission 490 nm).

### Powder X-ray diffraction

Purified Dysferlin C2A^WT^, C2A^W52R^ or C2A^V67D^ was suspended in 50 mM HEPES, 100 mM NaCl, pH 7.4. The insoluble fraction was centrifuged at 5000g for 10 min to generate a pellet. The pellet was resuspended in 1 ml 5 mM ammonium acetate followed by centrifugation at 5000g to remove residual HEPES/NaCl. The final pellet was resuspended in 20 µl of 5 mM ammonium acetate, pH 7.4, and pulled into a 0.7 mm quartz capillary tube. The sample was allowed to air dry in the presence of a desiccant. At 5 mM ammonium acetate is a volatile buffer and does not form crystals that would interfere with the diffraction pattern. X-ray powder diffraction data were acquired using a Rigaku Screen Machine (Rigaku) X-ray generator (50 kV, 0.6 mA) utilizing CuKα radiation (1.5418 °A) and a Mercury CCD detector. The distance from the sample to the detector was 75 mm. The samples were exposed to X-rays for 45 minutes.

### Molecular dynamics

Simulations for C2A^WT^ (4IHB), C2A^W52R^ and C2A^V67D^ were prepared in CHARMM-GUI. Mutations were introduced to the system using tools from CHARMM-GUI [28, 36]. Molecular dynamics simulations were performed using NAMD [46]. Simulations were performed using the CHARMM36m force field. TIP3 water model configuration was used throughout. Na^+^ and Cl^-^ were added at 150 mM to an electrically neutralized simulation box. Electrostatic forces were calculated using Particle-Mesh Ewald method (PME). A cut-off of 1.0 nm was used for calculating electrostatic and Van der Waals forces. Periodic boundary conditions were used. A short 100 ps NVT equilibration was initially performed. The velocity Verlet integrator MD simulation algorithm was implemented with a timestep of 2 fs. Coordinates were saved every 10 ps. The final production trajectory files were concatenated using PRODY CATDCD [65]. The MD production was run for 1000 ns using an NPT ensemble (Figure S3). H-bond analysis was performed with VMD (Figure 8).

### Transfection

HeLa cells were plated at 2 × 10^5^ cells per well of a 6-well plate and transfected with 750 ng of peGFP-N1, GFP-dysferlin, GFP-dysferlin^V67D^, or GFP-dysferlin^W52R^ using Lipofectamine2000 in Opti-MEM two days before cytotoxicity assays (Figure S4D).

Media was replaced one day after transfection. Transfection efficiencies for each construct ranged between 40-70% for each experiment (Figure S4D).

### Cytotoxicity Assay

Cytotoxicity assays were performed as previously described [21, 48, 51]. Briefly, 1×10^5^ cells were challenged in suspension with various concentrations of toxins for 30 min at 37 °C in RPMI supplemented with 2 mM CaCl_2_ (RC) and 20µg/mL propidium iodide (PI). Cells were analyzed on a 4-laser Attune Nxt flow cytometer (Thermo-Fisher). For analysis of cell lysis, we gated out the debris and then quantified the percentage of cells with high PI fluorescence (2–3 log shift). Prior work shows this PI^high^ population represents dead cells [31]. We calculated specific lysis as: % Specific Lysis = (% PI High^Experimental^ *−* % PI High^Control^)/ (100 *−* % PI High^Control^) × 100. The toxin dose needed to kill 50% of cells was defined as the Lethal Concentration 50% (LC_50_). LC_50_ was determined by logistic modeling using Excel (Microsoft, Redmond, WA, USA) as previously described [21].

### Inflammasome Assay

Dysferlin C2A^W52R^ or C2A^V67D^ was incubated overnight at 37°C prior to adding to macrophages. C2A^WT^ was similarly incubated overnight at 37 °C. 2.5 × 10^5^ BMDM were plated on 24-well plates and incubated at 37°C 5% CO_2_ overnight. Cells were primed with 100 EU/mL LPS for 4 h in Iscove’s modified Dulbecco’s medium. After washing, cells were incubated with no inhibitor, 50 mM KCl or 10 µM MNS for 30 min then stimulated with buffer, 20 µM nigericin for 30 min, 0.5 mg/mL MSU or 50 µg/mL Dysferlin wild-type C2A or 50 µg/mL Dysferlin mutant C2A for 1 h. Supernatants were collected and assayed either for pyroptosis by LDH assay or for IL-1β release by ELISA. Similarly, WT B6, NLRP3^-/-^, or Caspase 1/11^-/-^ BMDM were primed for 4 h with 100 EU/mL LPS, washed and stimulated with buffer, 20 µM nigericin for 30 min, 0.5 mg/mL MSU or 50 µg/mL C2A^WT^ or 50 µg/mL C2A^V67D^ for 1 h and supernatants collected and assayed for IL-1β production and LDH assay. IL-1β ELISA was performed as previously described [56], but using 2 µg/mL capture and 3 µg/mL detection antibody. OD values were read at 450 nm with 570 nm background subtraction using a Powerwave Microplate Spectrophotometer running Gen5 Data Analysis Software (BioTek, Winooski, VT). LDH assay was performed according to the manufacturer’s protocol, as previously described [24]. Supernatants were assayed in duplicate using the Pierce LDH cytotoxicity kit, using 1% Triton X-100 for 5 min as a positive control for complete cell lysis. OD was measured at 490 nm with 680 nm background subtraction using a Powerwave Microplate Spectrophotometer. Specific LDH release was determined as (mean OD value of sample *−* mean OD value of blank)/(mean OD value of Triton X-100 control sample *−* mean OD of blank) × 100%.

### Statistics

Prism 9.3.0 (GraphPad, San Diego, CA, USA), Origin (Northampton, MA) or Excel (Microsoft, Redmond, WA, USA) were used for statistical analysis. Data are represented as mean *±*SD as indicated. The LC_50_ for toxins was calculated by logistic modeling. Statistical significance was determined by either one-way ANOVA with Bonferroni post-testing or by repeated-measures ANOVA. p *<* 0.05 was considered to be statistically significant.

## Results

### C2A point-mutations in full-length dysferlin interfere with membrane repair

The pathogenic missense mutations C2A^W52R^ and C2A^V67D^ were tested for their ability to alter the capacity of full-length dysferlin protein to promote membrane repair. To test the consequences of these pathogenic point mutations on the activity of full-length dysferlin in a cell, we induced membrane damage using pore-forming toxins to stimulate the dysferlin repair pathway. Upon pore-formation, these toxins allow Ca^2+^ influx, which triggers dysferlin-mediated membrane repair [58]. The membrane repair potential of dysferlin was quantitated as a function of improved cellular resistance to toxin-induced cell death. We challenged transfected HeLa cells containing GFP alone, GFP-dysferlin^WT^, GFP-dysferlin^W52R^ or GFP-dysferlin^V67D^ with either aerolysin or streptolysin O (SLO). An inactive form of aerolysin (Y221G) was included as a negative control (Figure S4C). Consistent with our prior results [58], GFP-dysferlin^WT^ increased cellular resistance to toxin-induced cell death in HeLa cells *∼*4 fold when challenged with aerolysin (Figures 2A, S4A), and *∼*2 fold when challenged with SLO compared to controls (Figures 2B, S4B). Therefore, in comparison to GFP-dysferlin^WT^, GFP-dysferlin^W52R^ demonstrated a significant reduction in resistance to toxin-induced cell death, whereas GFP-dysferlin^V67D^ showed an overall trend toward reduced resistance. Thus, a single amino acid substitution in one of the eight C2 domains was sufficient to reduce dysferlin’s membrane repair capacity.

**Figure 2.**
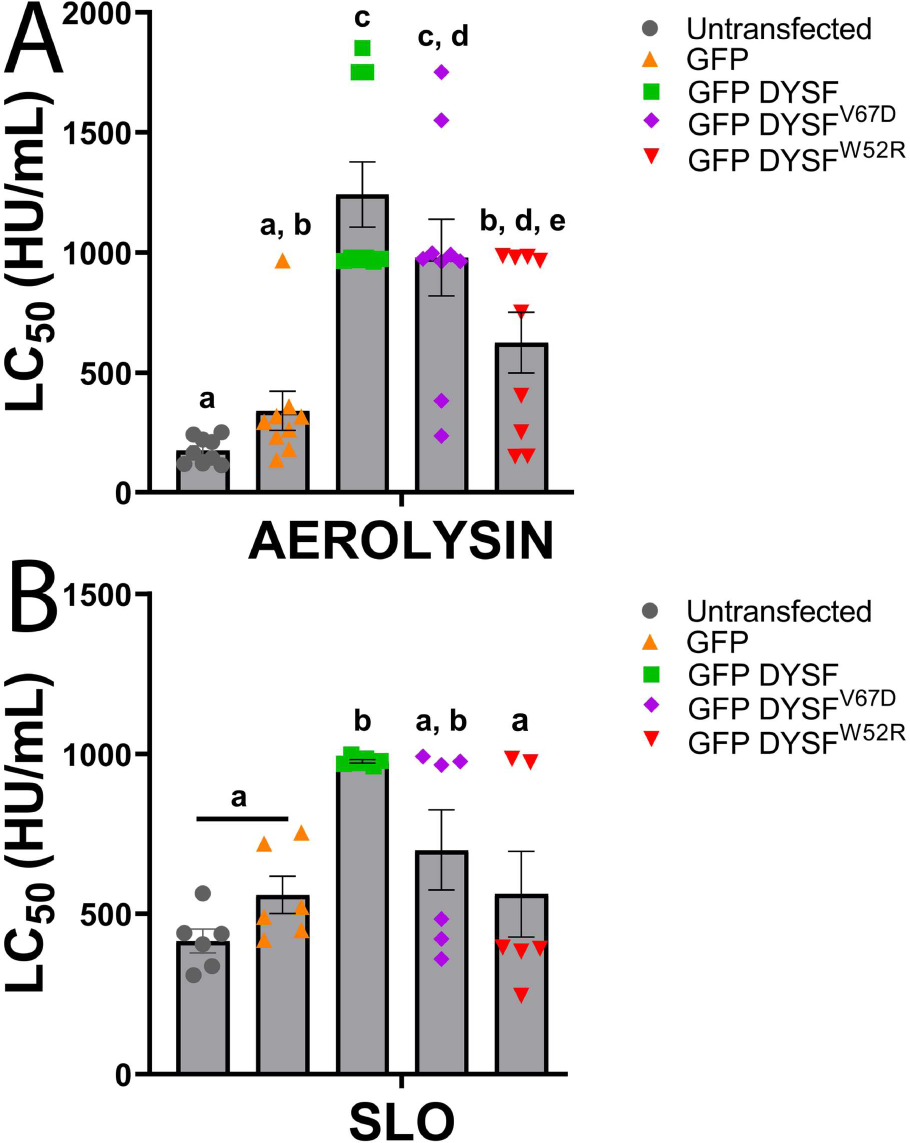
Mutations in the C2A domain of dysferlin reduce the protective activity of dysferlin. HeLa cells were transfected with GFP, GFP-dysferlin^WT^, GFP-dysferlin^V67D^, or GFP-dysferlin^W52R^ then incubated for 48 h and challenged with 31-2000 HU/mL of (A) aerolysin or (B) SLO in 2 mM CaCl_2_ supplemented RPMI including 20 µg/mL propidium iodide (PI) for 30 min at 37 °C. PI uptake was analyzed by flow cytometry to determine specific cell lysis. The amount of toxin needed to kill 50% of the cells (LC_50_) at each time point was calculated by logistic regression. Graphs display the average LC_50_ *±*S.E.M of (A) 9 or (B) 6 independent experiments. Letters (a-e) denote statistically significant (p *<* 0.05) groups using repeated-measures ANOVA between groups.

### Pathogenic point mutations destabilize the 3D structure of C2A

From an alignment of 3689 protein sequences annotated as dysferlin, the Trp-52 and the Val-67 positions are well conserved (Figure S5). The Trp-52 position occurs in the alignment as either a Trp or a Phe, emphasizing the bulky hydrophobic requirement for this locus. The most common variant for Val-67 in the dysferlin alignment is an Ile residue. Therefore, mutations that code for polar amino acids such as Arg and Asp would likely be deleterious to the fold of the domain. To determine whether the mutant C2A domains share secondary structure with C2A^WT^, we first determined the secondary structure content of purified C2A^WT^, C2A^W52R^, and C2A^V67D^ with and without Ca^2+^ by circular dichroism (CD) (Figure S2, Table 2). For C2A^WT^, we measured approximately 40% β-strand (Table 2). According to PDBMD2CD [13], the X-ray crystal structure of dysferlin C2A (PDB code 4IHB) has approximately 46% anti-parallel β-strand and a small fraction of α-helix. In both the C2A^W52R^ and C2A^V67D^ domains, we measured less total β-strand and more unordered peptide compared to C2A^WT^. We conclude that the high proportion of unordered structure, with or without Ca^2+^, is consistent with domain destabilization caused by mutations.

**Table 2.** Summary of the secondary structure content of each of the purified C2 domains in this study. Freshly purified, soluble C2 domain was assessed by CD prior to the triggering of amyloid by the removal of Ca^2+^ by EGTA. Secondary structure quantitation was evaluated using BestSel with a 200 - 250 nm data range. The NRMSD is a goodness-of-fit parameter that quantifies the difference between the experimental CD and computed spectra [43].

| Sample | Total Helix (%) | Total Strand (%) | Turns (%) | Unordered (%) | NRMSD |
| --- | --- | --- | --- | --- | --- |
| C2A<sup>WT</sup>+Ca | 7.2 | 38.7 | 14.3 | 39.7 | 0.01387 |
| C2A<sup>WT</sup>-Ca | 5.8 | 39.6 | 14.9 | 39.7 | 0.01759 |
| C2A<sup>W52R</sup>+Ca | 1.6 | 28.0 | 17.0 | 53.4 | 0.03177 |
| C2A<sup>W52R</sup>-Ca | 3.0 | 34.8 | 14.9 | 47.2 | 0.02931 |
| C2A<sup>V67D</sup>+Ca | 5.8 | 30.9 | 13.9 | 49.4 | 0.02157 |
| C2A<sup>V67D</sup>-Ca | 5.7 | 30.9 | 14.0 | 49.5 | 0.01830 |

Tryptophan-52 and Valine-67 of dysferlin’s C2A domain are integral constituents of the domain’s hydrophobic core (Figure 1). In the dysferlin C2A^WT^ X-ray structure (4IHB), approximately 1% of the Trp-52 residue is exposed to solvent, while the Val-67 locus is completely occluded as calculated by the Shrake-Rupley solvent-accessible surface area algorithm [7]. To test for the loss of integrity in the hydrophobic core in the wild-type and mutant C2A domains, we used the fluorescent dye ANS (1,8 ANS (1-anilinonaphthalene-8-sulfonic acid). Increased ANS fluorescence typically correlates with the dye’s accessibility to a hydrophobic environment, while lower ANS fluorescence correlates with exposure of the dye to an aqueous environment [53]. In the case of C2A^WT^ with bound Ca^2+^, we measured negligible binding of ANS, indicative of a protected hydrophobic core with minimal solvent accessibility (Figure 3). When Ca^2+^ is removed, C2A^WT^ showed increased ANS fluorescence (Figure 3), consistent with the low thermodynamic stability measured for the Ca^2+^-free domain (*<*1 kcal/mol) [17, 62]. Conversely, the C2A^W52R^ and C2A^V67D^ show a larger increase in ANS fluorescence when compared to C2A^WT^, regardless of the C2A domain’s Ca^2+^ bound state (Figure The blue-shift of the fluorescence for the C2A^W52R^ and C2A^V67D^ experiments provides additional evidence that the dye binds to the domains in a hydrophobic environment [22]. We conclude that the pathogenic point mutations within the C2A domain of dysferlin destabilize the folded domain via an increase in the exposure of their hydrophobic cores.

**Figure 3.**
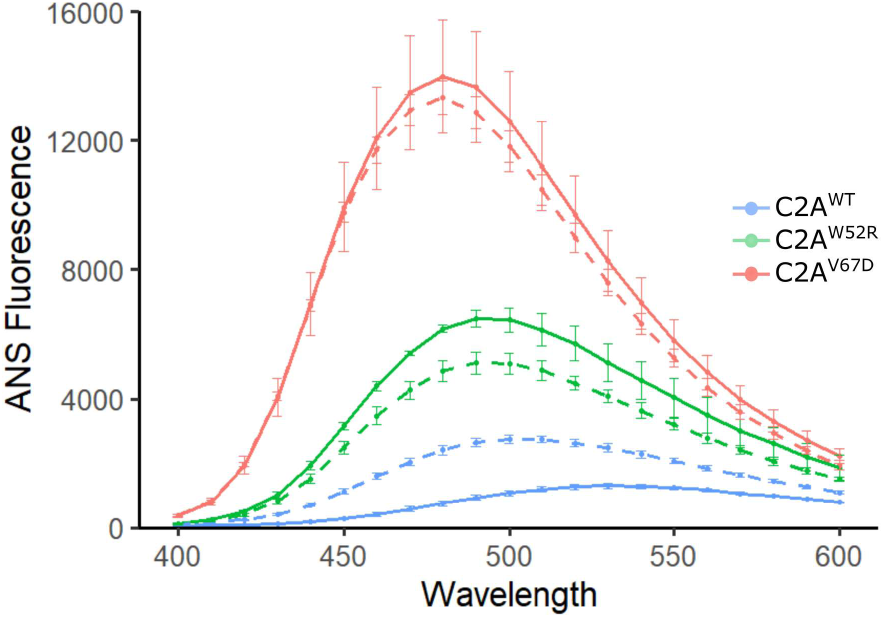
ANS binding was performed using purified dysferlin C2A, which was incubated with ANS dye. Each experiment was repeated with an n = 4. Error bars correspond to standard deviations, Ca^2+^-treated samples are represented by solid lines and EGTA-treated samples are represented by dashed lines.

### Pathogenic point mutations in C2A domains induce amyloid formation

The two C2A mutations were more prone to *in vitro* aggregation than the C2A^WT^, which made them more challenging to purify. Amyloid formation is one potential cause for this behavior. To test for the formation of dysferlin C2A amyloid, we incubated purified C2A domains with Thioflavin-T (ThT), a fluorescent dye used to test for the assembly of amyloid structures [44]. The C2A^WT^ domain exhibited low ThT fluorescence, with or without added Ca^2+^, indicating a lack of favorable binding sites for the dye and a low tendency for amyloid formation under our experimental conditions (Figure 4). C2A^W52R^ displayed an *∼*5-fold increase in ThT fluorescence compared to the C2A^WT^. C2A^V67D^ showed the highest tendency to bind ThT at*∼* 7-10-fold higher fluorescence signal than the C2A^WT^. Interestingly, the Ca^2+^-bound form of C2A^V67D^ was more prone to amyloid formation than the Ca^2+^-free form (Figure 4), implying Ca^2+^-dependent instability of the C2A^V67D^ domain. In contrast, the C2A^W52R^ mutation was less likely to generate amyloid in the presence of Ca^2+^ than its Ca^2+^-free version. These findings are consistent with Ca^2+^-induced domain stability, which has been noted in other C2 domains [55].

**Figure 4.**
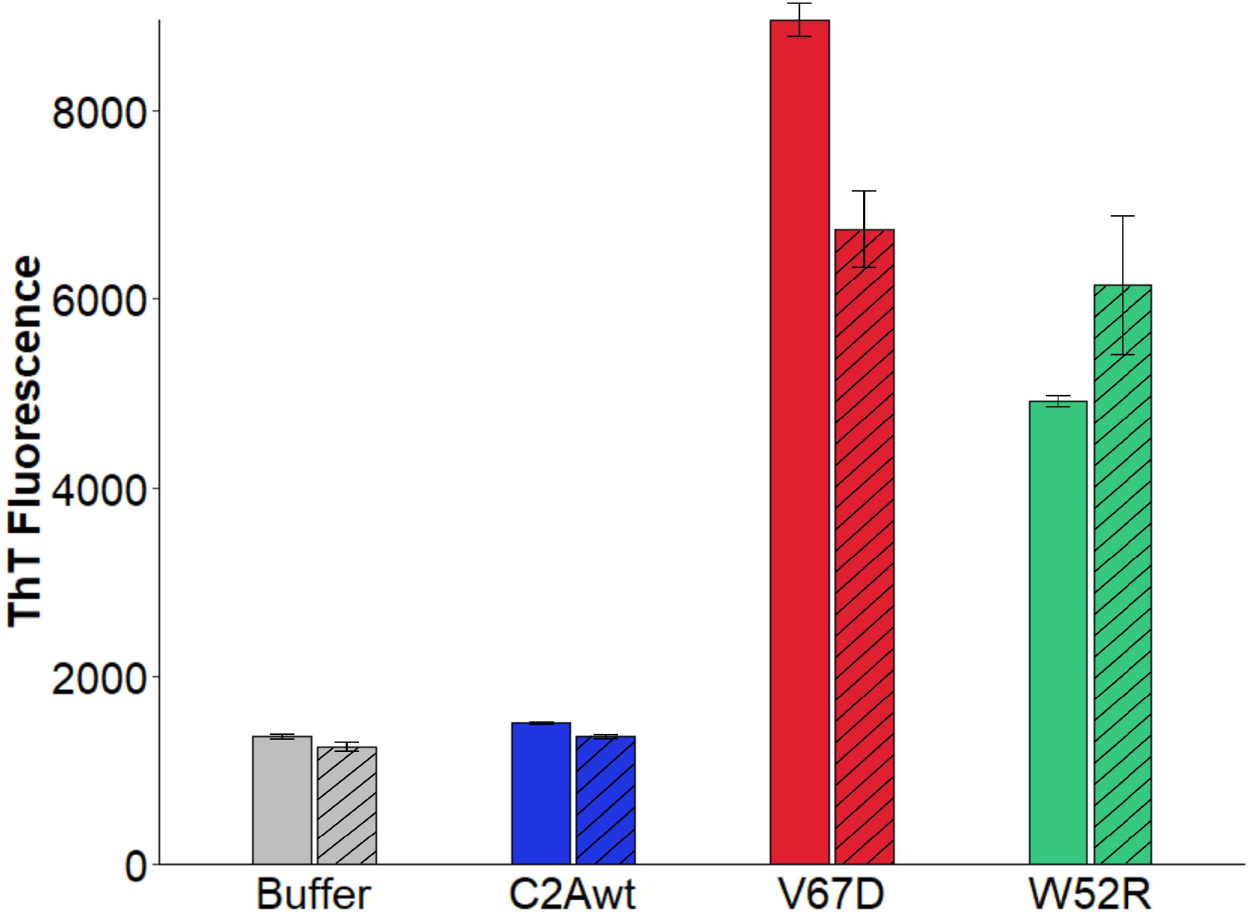
Thioflavin-T binding to purified C2A domain. Solid-colored bars correspond to Ca^2+^-treated C2A domains, while hashed bars correspond to EGTA-treated samples (n=3). The chart displays error bars as standard deviation.

To verify that mutant C2A domains can form amyloid structures, we used X-ray fiber diffraction on the purified, mutant dysferlin C2A domains. Fiber diffraction has traditionally been used to characterize periodic geometries in aligned or unaligned amyloid samples [54]. C2A^WT^ did not form a solid under our experimental conditions and consequently could not be measured by this method. Consistent with measurements of other amyloids, our analysis yielded two primary reflections characteristic of cross-β structure [54]. For the C2A^W52R^ sample, we measured a 4.6 °A meridional reflection and a 10.2 °A equatorial reflection (Figure 5). For the C2A^V67D^ sample, we measured a meridional reflection at 4.4 °A and an equatorial reflection at 9.8 °A. The meridional reflection suggests a regular structural repeat of *∼*4 °A along the axis of the structure; the equatorial reflection reveals a structural spacing of *∼*10 °A perpendicular to the axis of the structure. We conclude that the material prepared using purified mutant dysferlin C2A domains possesses cross-β sheet structure similar to that found in other amyloids.

**Figure 5.**
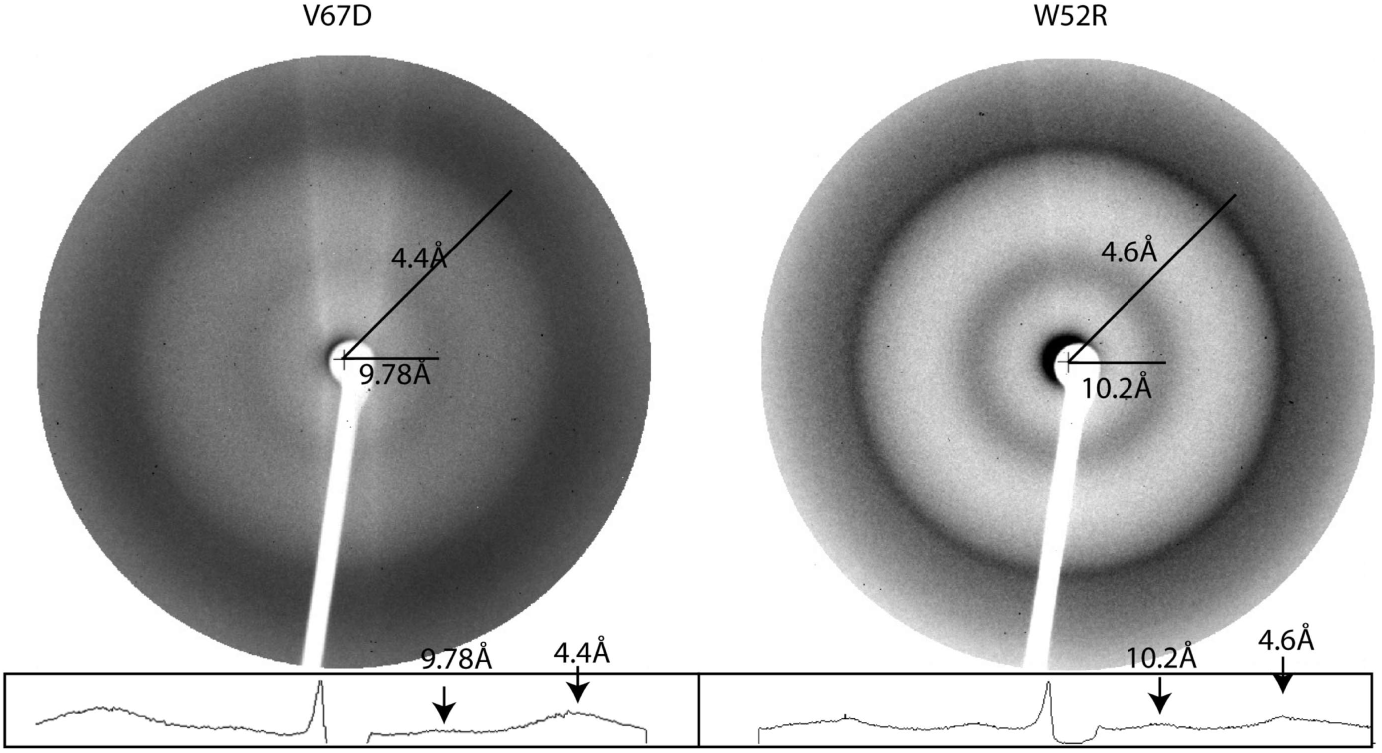
Un-oriented fiber diffraction of dysferlin C2A^W52R^ and dysferlin C2A^V67D^ amyloid material. The radial averaging exhibits reflections associated with inter- and intra-β-sheet spacings consistent with amyloid structures. The line graph below the diffraction image is the radial cross-section of intensity across the center of the X-ray image. The relevant maxima are indicated by arrows and their corresponding distances from the center of the image.

We have confirmed that single amino acid mutations among the C2 domains of dysferlin are sufficient to cause dysfunction in the full-length dysferlin protein (Figure 2). Further, the point mutations in the isolated C2 domains result in domain misfolding, which forms material consistent with classical amyloid structure *in vitro*. To assess the specific locus within the dysferlin C2A sequence most capable of nucleating amyloid, we analyzed the primary sequence of the dysferlin C2A domain using Amylpred2. Amylpred2 combines the results of multiple amyloid assessment algorithms to score a protein’s primary sequence according to its amyloid-forming propensity [61]. According to this analysis of dysferlin C2A^WT^, amyloid-prone β-strands correspond to β-strands 1, 2, 5, and 8 (Figure 6B). In the crystal structure of the C2A^WT^ domain, β-strands 1 and 8 are hydrogen bonded in an anti-parallel β-sheet configuration, constituting the two innermost strands of sheet A (Figure 1. Similarly, β-strands 2 and 5 hydrogen bond to establish the two innermost strands of sheet B (Figure 1).

**Figure 6.**
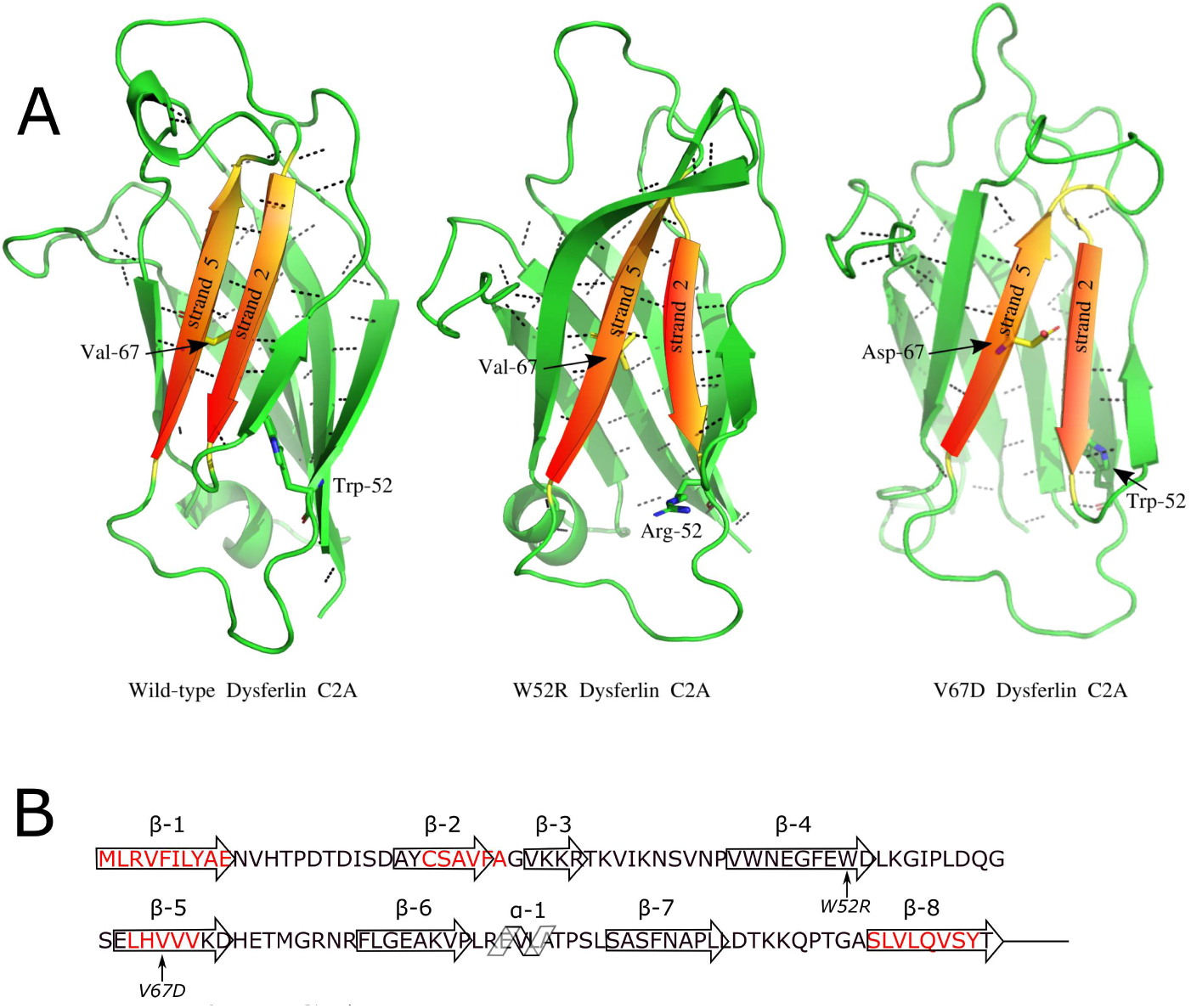
**A.** Dysferlin C2A mutations increase the domain’s amyloidogenic potential. Snapshots of MD sims from Dysferlin C2A^WT,^ C2A^W52R^ and C2A^6V7D^ are shown. Dashed lines correspond to backbone H-bonds at least 3.0 Å and 20° from donor to acceptor. Shaded β-strands correspond to the magnitude of the amyloid-forming propensity of the sequences exposed as a function of the mutations, as measured by Amylpred. **B.** Primary sequence of human dysferlin C2A. The secondary structure elements of the C2 domain are superimposed on the sequence. Arrows represent β-strands, helix α-1 is shown as a helical schematic. Residues in red correspond to positive matches for amyloidogenic sequences according to Amylpred. The mutated residues W52R and V67D are indicated with arrows.

Thus Amylprep analysis indicates that the two central β-strands in each of the two β-sheets show a strong propensity to form amyloid (Figure 6B); however, in the wild-type C2A domain fold, β-stands 2 and 5 are shielded by flanking β-strands 3 and 6. It is unclear how these β-strands are exposed to form an amyloid structure. To determine a rationale for the structural defect caused by the two missense mutations in dysferlin C2A, we ran 1 µs molecular dynamics (MD) simulations of C2A^WT^, C2A^W52R^, and C2A^V67D^. In the dysferlin C2A^W52R^ simulation, the most obvious change in the structure was a widening rupture in the space between β-strands 2 and 5 on sheet B (Figure 6A). Initially, the gap in sheet B was a puzzling result since the Trp-52 residue is on the opposite sheet from the rupture. Over the course of the simulation, the rupture continued to widen, as indicated by the gradual loss of the backbone H-bonds between β-stands 2 and 5 (Figure S4).

In the dysferlin C2A^V67D^ simulation, Charmm-GUI modeled the polar Asp-67 mutation in the same orientation as the buried Val-67 residue. As the simulation progressed, Asp-67 quickly migrated from its initial internal configuration to a solvent-exposed orientation. Backbone H-bonds joining the anti-parallel β-sheet were disrupted to accommodate this structural shift. Therefore, we predict that the V67D mutation disrupts the backbone H-bonds that hold β-sheet ‘B’ together (Figures 1, 6). Hbond analysis of the three trajectories shows that C2A^WT^ maintains an average of eight H-bonds between strands 2 and 5, along the length of sheet ‘B’ (Figure S4). This contrasts with the C2A^W52R^ case, where an average of 3 backbone H-bonds formed between β-stands 2 and 5, and an average of 4 backbone H-bonds for C2A^V67D^ (Figure S4).

### Electron micrographs of C2A show structures consistent with other non-peptide amyloid

We next studied the appearance of C2A^W52R^ and the C2A^V67D^ amyloids by negative stained, Transmission Electron Microscopy. In the TEM micrographs, C2A^WT^ formed very little structure, while the C2A^W52R^ and the C2A^V67D^ formed film-like structures consistent with the appearance of other known non-peptide amyloids (Figure S6). The C2A amyloid that we imaged was similar in appearance to the amyloid structures derived from full-length proteins observed for *Streptococcus mutans* [3] and Cystatin-Related Epididymal Spermatogenic (CRES) [11].

### Dysferlin C2A mutations activate the NLRP3 inflammasome

Next, we evaluated the C2A’s potential to induce inflammation. Activation of the Nod-Like Receptor family, pyrin domain containing 3 (NLRP3) inflammasome is an important pro-inflammatory mechanism. [30]. Given that amyloid-β can activate NLRP3 [19], we tested whether the NLRP3 inflammasome could be activated by the dysferlin C2A amyloid produced from C2A^W52R^ or C2A^V67D^. To test whether these dysferlin amyloids trigger inflammasome activation, LPS-primed bone marrow-derived macrophages (BMDM) were treated with either the positive controls, nigericin and monosodium urate (MSU) crystals, or C2A^WT^, C2A^W52R^ or C2A^V67D^. The purified C2A^WT^ sample controlled both the inherent inflammasome activation potential of dysferlin C2A domains and the presence of other contaminating NOD-like receptor (NLR) agonists in the purified protein preparation. Nigericin, a K^+^ ionophore, induced robust IL-1β secretion (Figure 7a). MSU triggers the NLRP3 inflammasome via lysosomal damage in a manner consistent with amyloids [38]. MSU triggered robust IL-1β secretion (Figure 7a). Compared to MSU, C2A^W52R^ and C2A^V67D^ induced similar levels of IL-1β secretion in LPS-primed BMDM, whereas C2A^WT^ failed to do so (Figure 7a). This suggests that C2A^W52R^ and C2A^V67D^ amyloids can stimulate the release of pro-inflammatory cytokines. To test if NLRP3 senses the C2A^W52R^ and C2A^V67D^ amyloids, we used the specific NLRP3 inhibitors KCl [45] or 3,4-methylenedioxy-β-nitrostyrene (MNS) [23]. We found that both KCl and MNS inhibited IL-1β release from BMDM (Figure 7a). These data suggest C2A^W52R^ and C2A^V67D^ amyloid trigger the NLRP3 inflammasome. To confirm that C2A^W52R^ and C2A^V67D^ amyloid act via an NLRP3-dependent mechanism, we treated LPS-primed B6 or NLRP3^-/-^ BMDM with nigericin, MSU, C2A^WT^, C2A^W52R^ or C2A^V67D^. In contrast to B6 BMDM, NLRP3^-/-^ BMDM did not produce IL-1β in response to any of the stimuli (Figure 7b). To test if the inflammatory caspases, Caspase-1 or Caspase-11, are required, we treated Casp1/11^-/-^ BMDM with NLRP3 agonists, dysferlin C2A^WT^, dysferlin C2A^W52R^ amyloid or dysferlin C2A^V67D^ amyloid. We found that IL-1β release was blocked in Casp1/11^-/-^ BMDM following treatment with nigericin, MSU, or dysferlin C2A^W52R^ and dysferlin C2A^V67D^ amyloid (Figure7b). Thus, these results suggest that dysferlin C2A^W52R^ and dysferlin C2A^V67D^ amyloid promote NLRP3 inflammasome-dependent IL-1β release.

**Figure 7.**
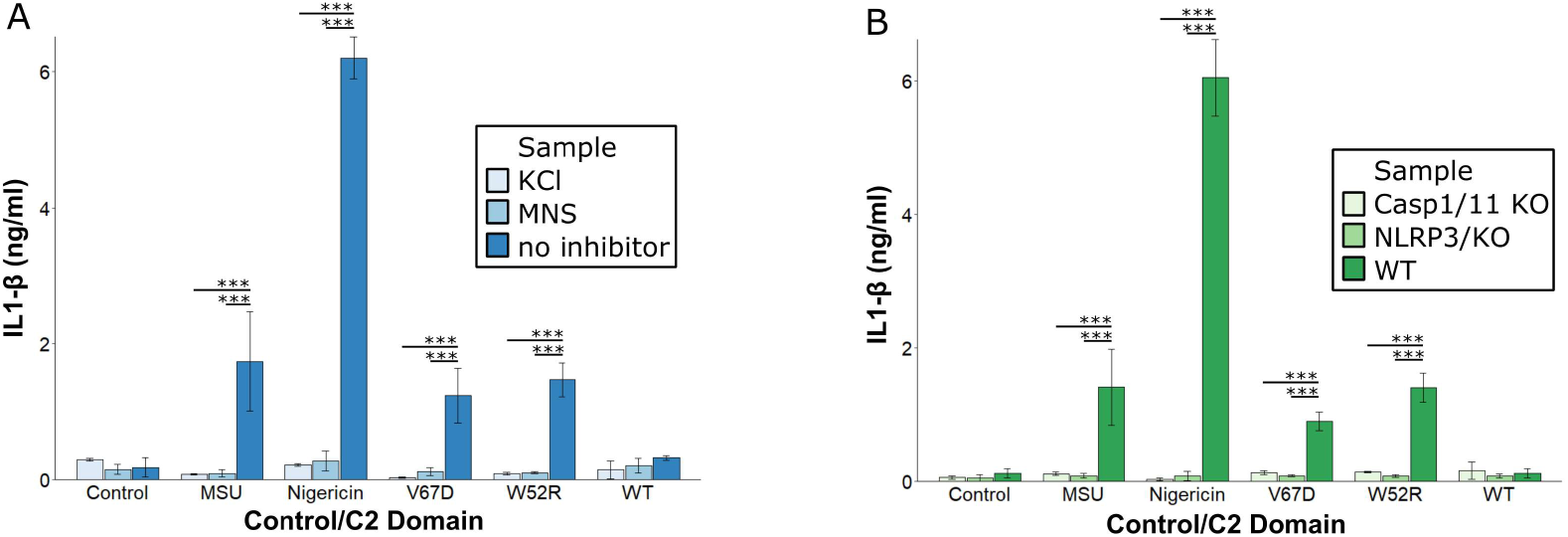
Dysferlin C2A pathogenic point mutants induces NLRP3-dependent IL-β release. (A) B6 BMDM were primed with 100 EU /mL LPS for 4 hrs, washed, treated with no inhibitor, 50 mM KCl or 10 ¼M MNS for 30 min, and stimulated with nothing, 20 ¼M nigericin for 30 min, 0.5 mg/mL MSU, 50 ¼g/mL C2A^WT^ or 50 ¼g/mL C2A^V67D^ for 1 h. Supernatants were collected and assayed for IL-β1 production by ELISA. (B) B6, NLRP3^-/-^ or Caspase 1/11^-/-^ BMDM were primed with 100 EU /mL LPS for 4 h, washed and stimulated with nothing, 20 ¼M nigericin for 30 min, 0.5 mg/mL MSU, 50 ¼g/mL C2A^WT^ or 50 ¼g/mL C2A^W52R^ or C2A^V67D^ for 1 h. Supernatants were collected and assayed for IL-β production by ELISA. Graphs represent mean ±SEM of three independent experiments. *** p<0.001.

### Dysferlin C2A mutations induce pyroptosis

Another functional hallmark of inflammasome activation is Casp1/11-dependent programmed cell death, termed pyroptosis [5, 42]. To test whether C2A^W52R^ or C2A^V67D^ can trigger pyroptosis, we treated LPS-primed BMDM with nigericin, MSU, dysferlin C2A^WT^, dysferlin C2A^W52R^ or dysferlin C2A^V67D^, and measured cell death by lactate dehydrogenase (LDH) release, as previously done [24]. Using this assay, nigericin and MSU induced significant cytotoxicity (Figure 8a). Dysferlin C2A^W52R^ and dysferlin C2A^V67D^ also induced cell death in LPS-primed BMDM, whereas dysferlin C2A^WT^ failed to do so (Figure 8a). To determine if dysferlin C2A^W52R^ and dysferlin C2A^V67D^ amyloid-induced cell death required NLRP3, we used the NLRP3-specific inhibitors KCl and MNS. We found that either KCl or MNS reduced BMDM cell death following nigericin, MSU, dysferlin C2A^W52R^ or dysferlin C2A^V67D^ treatment (Figure 8b). We next confirmed the specificity of these inhibitors using knockout mice. LPS-primed NLRP3^-/-^ BMDM did not show cell death, in contrast to wild-type BMDM (Figure 8b). Therefore our data indicate that dysferlin C2A^W52R^ or dysferlin C2A^V67D^ mediated cell death via the NLRP3 inflammasome. Since one key hallmark of pyroptosis is Casp1/11 dependence, we tested whether dysferlin C2A^W52R^ and dysferlin C2A^V67D^ could induce cell death in Casp1/11^-/-^ BMDM. We found that cell death induced by dysferlin C2A^W52R^ and dysferlin C2A^V67D^, nigericin, or MSU was abrogated in Casp1/11^-/-^ BMDM (Figure 8b). Based on these data, we conclude that dysferlin C2A^W52R^ and dysferlin C2A^V67D^ activate the NLRP3 inflammasome to induce pyroptosis in macrophages. Altogether, our results suggest that amyloid-induced inflammasome activation may underlie the inflammatory pathogenesis of Limb-Girdle Muscular Dystrophy Type 2B and Miyoshi Myopathy.

**Figure 8.**
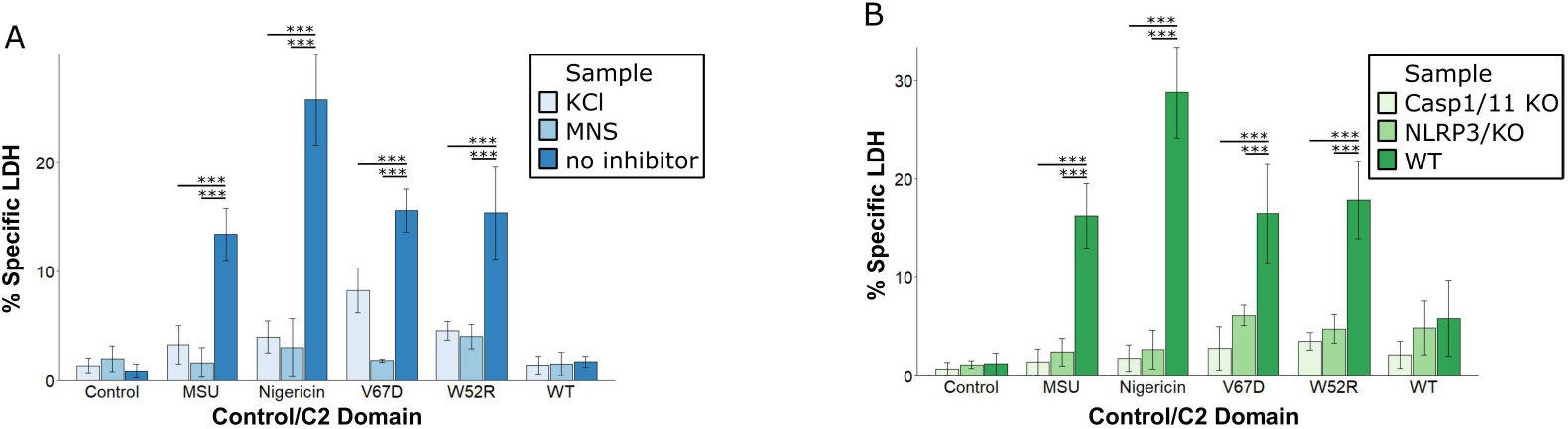
LPS-primed BMDM were treated as described in Figure (7) and supernatants assayed for LDH release. LPS-primed BMDM lysed with 1% Triton were used for maximum release. Graphs represent mean ±SD of three independent experiments. * p<0.05, ** p<0.01, *** p<0.001.

## Discussion

Here we found that pathogenic dysferlin mutations V67D and W52R induce dysferlin to form pro-inflammatory amyloids. These mutations disrupt the folding of the C2A domain and interfere with dysferlin function. X-ray scattering consistent with cross-β-amyloid formation and amyloid-specific ThT staining was observed in mutant dysferlin C2A domains. Additionally, the dysferlin C2A amyloids could activate the NLRP3 inflammasome. Together our data provide a mechanistic basis for inflammatory disease during dysferlinopathies.

Mutations in the *DYSF* gene are the underlying cause of Limb-Girdle Muscular Dystrophies Type-2B/2R and Miyoshi Myopathies. These mutations occur evenly throughout the gene, causing altered gene products, including truncated proteins and missense mutations within protein domains. To date, approximately 270 missense mutations associated with the disease have been described in the gene [6], but no genomic “hotspots” has yet been identified in *DYSF*. The most extensively studied mutations in the C2A domain of dysferlin are C2A^W52R^ and C2A^V67D^. Our results show that these mutations alter proper membrane localization and impair membrane repair, as cells with either pathogenic mutation were unable to enhance repair compared to wild-type GFP-dysferlin [58]. These findings suggest that these single-point mutations in a single domain can compromise the entire protein’s function.

The postulated reason for dysfunction in the C2A domain of dysferlin is that the two mutations, C2A^W52R^ and C2A^V67D^, inadvertently expose amyloidogenic regions that are normally protected by the assembly of the β-sheets of the C2A domain. The mutations C2A^W52R^ and C2A^V67D^ result in the loss of hydrogen bonds in sheet B of the C2A domain (Figure 8). The decreased loss of backbone H-bond sheet integrity is observed for both mutations, even though they occur on different sheets. The absence of the bulky Trp side chain could have created defects in the hydrophobic core of the C2A domain, leading to a cleft in the opposite sheet. In the dysferlin C2A^WT^, the hydrogen bonding throughout the length of the sheet conceals β-strands 2 and 5 from forming non-productive intra-domain interactions. The proposed hypothesis is that the two mutations in C2A cause these strands to misfold, exposing both the amyloidogenic sequences and the hydrophobic core, leading to the formation of amyloid. The formation of the cross-β structures and resulting amyloid was confirmed through ThT staining (Figure 4) and X-ray diffraction (Figure 5).

We investigated if dysferlin-based amyloids, like other pathogenic amyloids, promote inflammation. Our findings showed that both dysferlin C2A mutations activate the NLRP3 inflammasome, which is expressed in muscle cells [16] and known to be triggered by Amyloid-β [19]. This suggests that pathogenic amyloids could cause muscle degeneration through pyroptosis. Similarly, in a mouse model of Duchenne’s Muscular Dystrophy, treatment with an NLRP3 inhibitor reduced inflammation and disease severity, emphasizing the crucial role of inflammasome activation in the disease [14]. Our results indicate that amyloidogenesis could be the starting point for the inflammation observed during disease progression. Further research is needed to determine how other dysferlin mutations increase amyloidogenesis [52, 57].

The C2A^W52R^ and C2A^V67D^ mutations affect a dual negative impact on LGMD-2B/2R patients. Firstly, these mutations cause dysferlin to lose its normal function. In a patient muscle biopsy, W52R was found to be “absent” from the membrane [37], while only 16% of V67D was found to be localized to the membrane [59]. The C2A domain, as amyloid, can no longer sense Ca^2+^ or bind to phospholipids [10]. Instead, the domain forms an insoluble amyloid by exposing sequences prone to forming amyloids. The mutant C2A dysferlin partitions away from the membrane, thus reducing its ability to contribute to repair. Finally, the amyloid could lead to the death of muscle cells via pyroptosis, releasing signals that trigger an immune response. The resulting inflammation could then be accelerated by IL-1β produced by macrophages.

## Acknowledgments

Research reported in this publication was, in part, supported by the Take Part Foundation (https://take-part.org/), the National Institute of Arthritis and Musculoskeletal and Skin Diseases (R01 AR063634, to R.B.S), the National Institute of Allergy and Infectious Disease (R21 AI156225, to P.A.K), and American Heart Association (16SDG30200001, to P.A.K), and National Institute of Child Health and Human Development (R01 HD033903, to G.A.C). The authors would also like to acknowledge the Jain Foundation (https://www.jain-foundation.org/) for support of this research. The authors acknowledge the High-Performance Computing Center (HPCC) at Texas Tech University at Lubbock (http://cmsdev.ttu.edu/hpcc) and Texas Advanced Computing Center (TACC) at The University of Texas at Austin (http://www.tacc.utexas.edu) for providing High-Performance Computing resources that have contributed to the research results reported within this paper. We thank the College of Arts & Sciences Microscopy at TTU for use of flow cytometry facilities.

