## Supplemental Information for "Pathogenic Mutations in the C2A Domain of Dysferlin form Amyloid that Activates the Inflammasome"

### Supporting Information

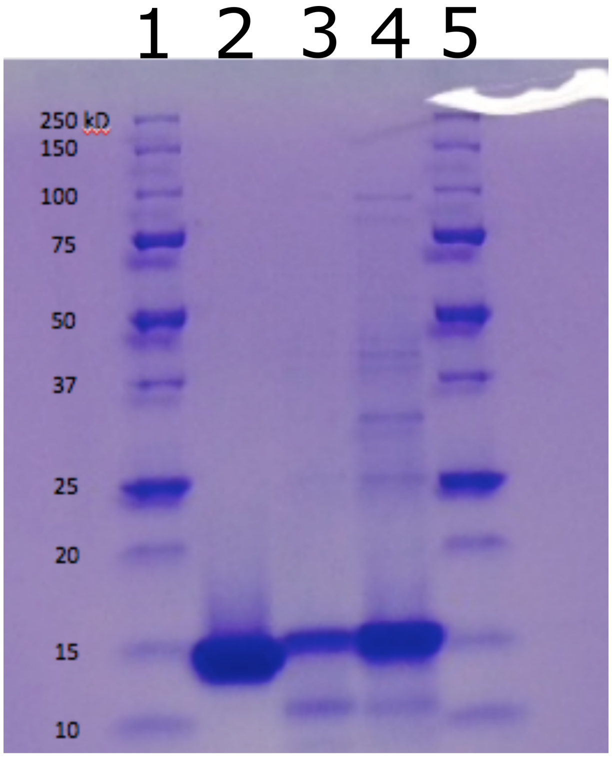


**Figure S1.** PhastGel of purified C2A$^{WT}$ (lane 2), C2A$^{W52R}$ (lane 3), and C2A$^{V67D}$ (lane 4). The gel was stained with Coomassie Blue. "Stuttering" of the loading comb produced the double loading appearance of some of the bands in lanes 1 and 5.

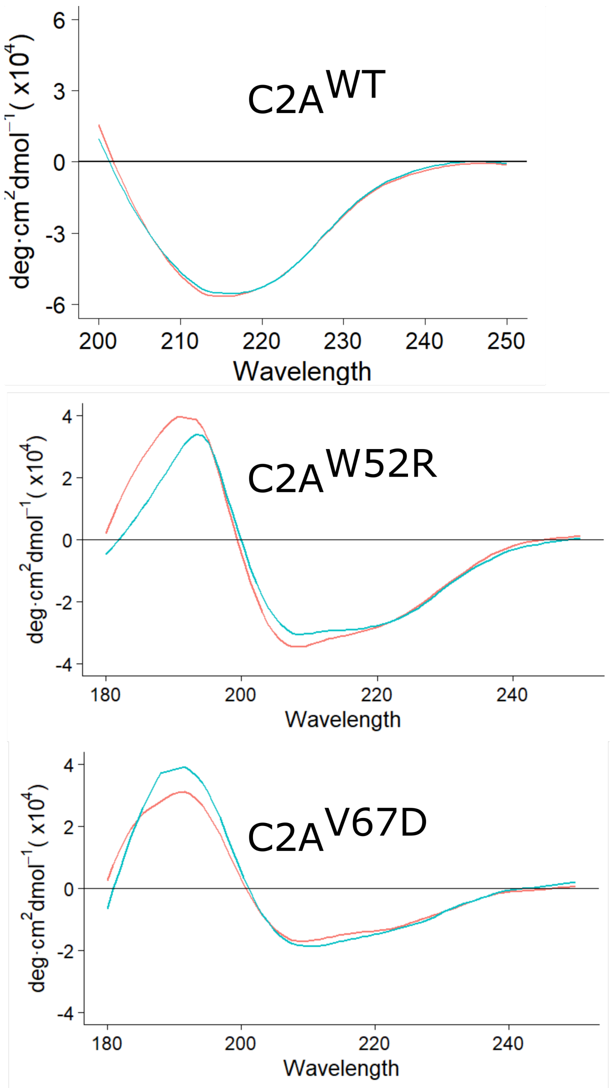


**Figure S2.** representative Circular Dichroism (CD) spectra of C2A$^{WT}$ (top), C2A$^{W52R}$ (middle), and C2A$^{V67D}$ (bottom). EGTA-treated, no $Ca^{2+}$ C2A domain is shown in red, while $Ca^{2+}$-treated domain is shown in green.

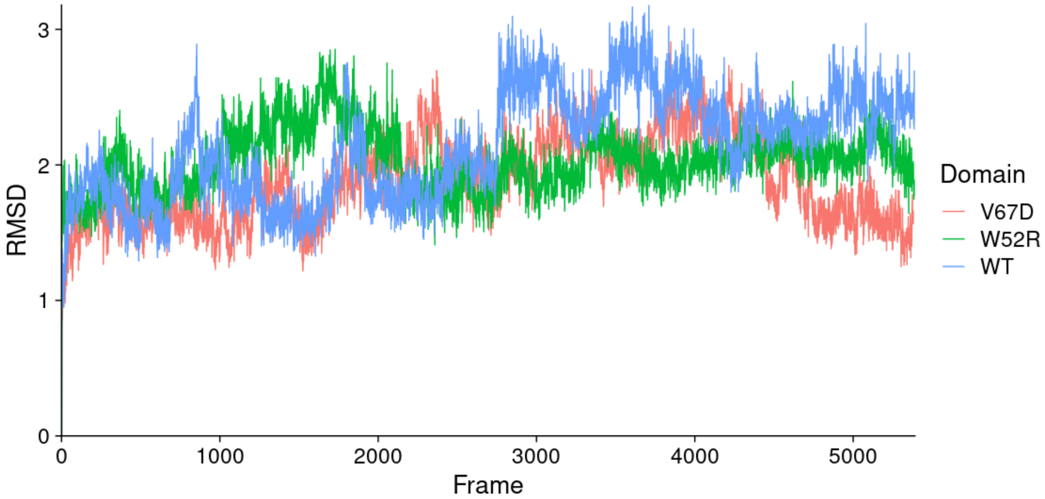


**Figure S3.** First 5000 frames of the C2A trajectories of dysferlin C2A$^{WT}$, C2A$^{W52R}$, and C2A$^{V67D}$. A total of 1250 ns of each domain was computed and analyzed.

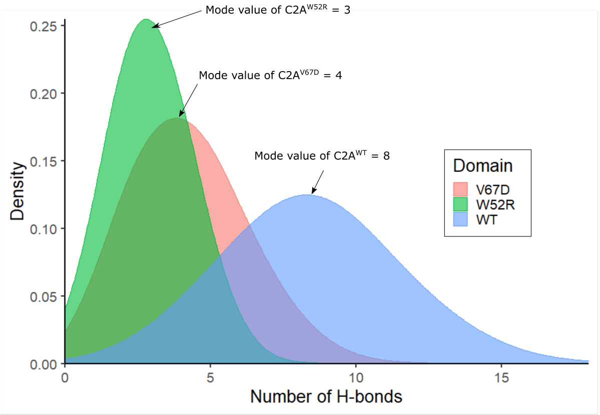


**Figure S4.** Histogram/Density plot showing the number of backbone H-bonds that form between β-strands 2 and 5 on sheet B over the time course of the molecular dynamics trajectory C2A$^{WT}$, C2A$^{W52R}$, and C2A$^{V67D}$. Hydrogen bonds were analyzed using the VMD Hbonds plugin. Only equilibrated trajectory frames were analyzed. The mode values of the various histograms represent the number of H-bonds that occurs most often in the data set.

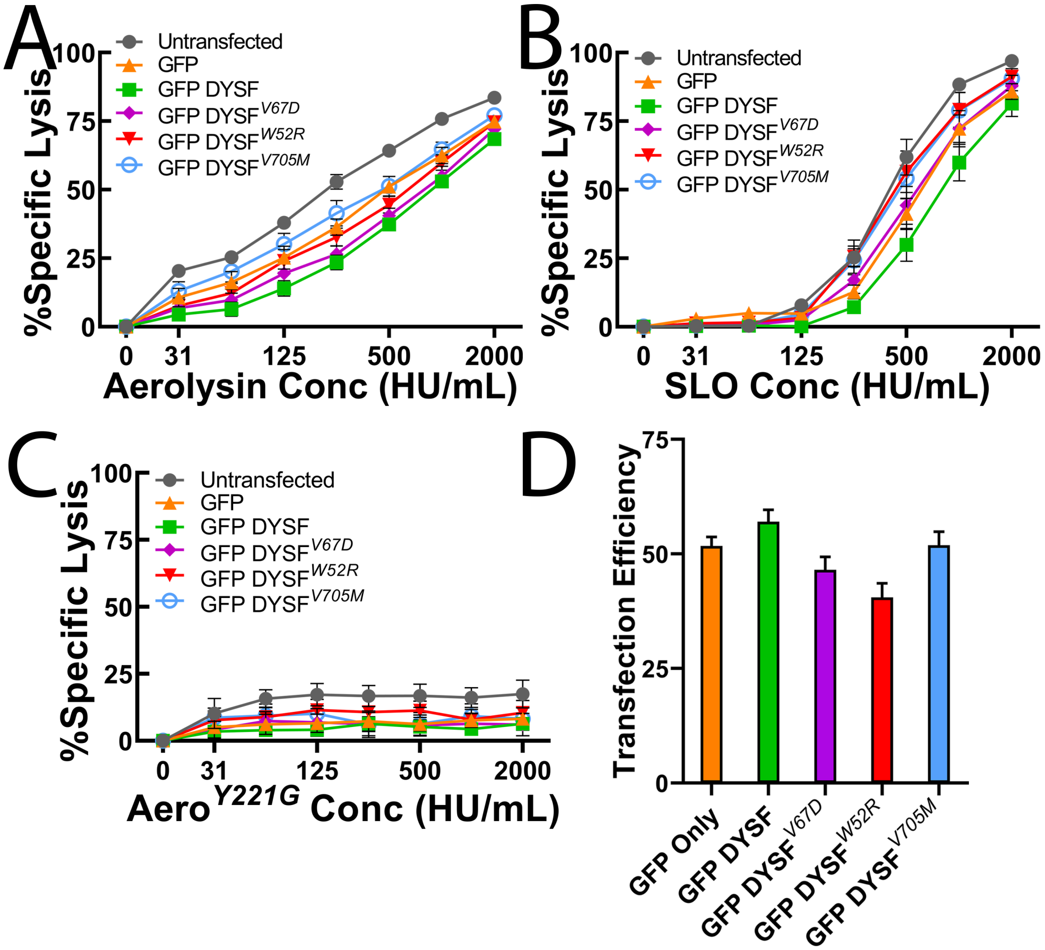


**Figure S5.** Mutations in dysferlin reduce dysferlin activity. HeLa cells were either untransfected or transfected with GFP, GFP-dysferlin DYSF, GFP- dysferlin$^{V67D}$, or GFP-dysferlin$^{W52R}$ for 48 h and challenged with 31-2000 HU/mL of (A) aerolysin, (B) SLO or (C) mass equivalent of mutant aerolysin$^{Y221G}$ in 2 mM $CaCl_2$ supplemented RPMI (RC) with 20 µg/ml propidium iodide (PI) for 30 min at 37°C. PI uptake was analyzed by flow cytometry and specific lysis was determined as described in Methods. (D) The transfection efficiency of transfected HeLa cells in A-C is shown. Graphs show the average ± S.E.M of (A, D) 9, (B) 6 or (C) 3 independent experiments.

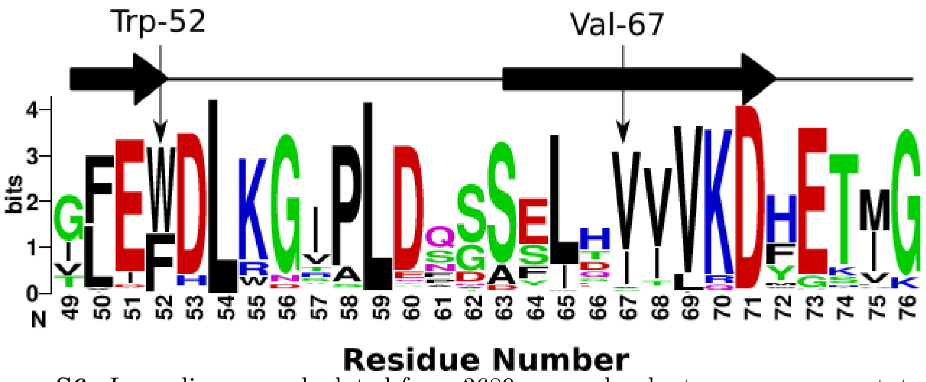


**Figure S6.** Logo diagram calculated from 3689 non-redundant sequences annotated as dysferlin. Residue numbering is human dysferlin. Residues 71 and 73 are $Ca^{2+}$ binding residues

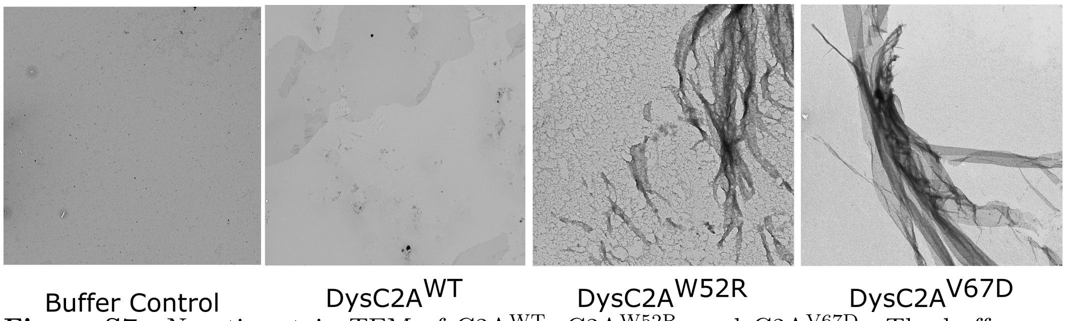


**Figure S7.** Negative stain TEM of C2A$^{WT}$, C2A$^{W52R}$, and C2A$^{V67D}$. The buffer control and the C2A$^{WT}$ are shown at 1000X direct magnification. C2A$^{W52R}$ micrograph is shown at 5000X. The C2A$^{V67D}$ is shown at 2000X direct magnification.
